# Canonical Wnt signaling regulates soft palate development through mediating ciliary homeostasis

**DOI:** 10.1101/2022.08.03.502715

**Authors:** Eva Janečková, Jifan Feng, Tingwei Guo, Xia Han, Siddhika Pareek, Aileen Ghobadi, Thach-Vu Ho, Angelita Araujo-Villalba, Jasmine Alvarez, Yang Chai

## Abstract

Craniofacial morphogenesis requires complex interactions among tissues, signaling pathways, secreted factors, and organelles. The details of these interactions remain elusive. In this study, we analyzed the molecular mechanisms and homeostatic cellular activities governing soft palate development to improve regenerative strategies for cleft palate patients. We have identified canonical Wnt signaling as a key signaling pathway primarily active in cranial neural crest (CNC)-derived mesenchymal cells surrounding soft palatal myogenic cells. Using *Osr2-Cre;β-catenin^fl/fl^* mice, we further discovered that Wnt signaling is indispensable for mesenchymal cell proliferation and subsequently myogenesis through mediating ciliogenesis. Specifically, we identified that Wnt signaling directly regulates expression of the ciliary gene *Ttll3* through β-catenin/Tcf7l2 complex. Impaired ciliary disassembly leads to differentiation defects of mesenchymal cells and indirectly disrupts myogenesis through decreased expression of *Dlk1*, a mesenchymal cell-derived pro-myogenesis factor. Moreover, we found that restoring ciliary homeostasis rescues mesenchymal cell proliferation in *Osr2-Cre;β-catenin^fl/fl^* samples. This study highlights the role of Wnt signaling in palatogenesis through controlling ciliary homeostasis, which establishes a new mechanism for Wnt-regulated craniofacial morphogenesis.

## Introduction

Organ morphogenesis is a complex process involving a combination of homeostatic cellular activities as well as reciprocal tissue-tissue interactions with signals relayed by secreted factors (Strand et al. 2010). The soft palate is an anatomical structure with unique functions, and its embryonic development involves interplay among cranial neural crest (CNC)-derived mesenchymal cells, mesoderm-derived myogenic cells, and ectoderm-derived epithelial cells, making it an excellent model for unraveling the intricate morphogenetic tissue-tissue communications during craniofacial development (Li et al. 2019, Grimaldi et al. 2015, Sugii et al. 2017). Of particular interest is the role of CNC-derived mesenchymal cells and the factors derived from these cells, which ultimately mediate the communication of these mesenchymal cells with the myogenic precursors during early craniofacial muscle development. The majority of these factors, mechanistic links, and the molecular crosstalk among them are yet to be discovered (Noden and Francis-West 2006, Sambasivan et al. 2011, Ziermann et al. 2018, Sugii et al. 2017).

Cleft lip with or without cleft palate, which occurs in 1/700 live births, is one of the most common birth defects and significantly affects the quality of life (Dixon et al. 2011). The mechanism associated with cleft soft palate is less understood than that of cleft hard palate, but soft palate clefting imposes a significant burden on patients and healthcare providers and may require treatment from infancy through adulthood (Wehby and Cassell 2010). The soft palate is part of a larger structure known as, the oropharyngeal complex, and its main functions are in swallowing, breathing, hearing, and speech, which are disrupted in patients with clefts. Due to the delicate structure of the soft palate, its functional restoration requires long-term and multidisciplinary treatment (Monroy et al. 2012). Soft palatal functions are not sufficiently re-established in 30% of cleft patients after surgery (Von den Hoff et al. 2018, Monroy et al. 2015, Marrinan et al. 1998).

To date, studies have identified *Dlx5*/*Fgf10*, *Mn1*/*Tbx22*, *Runx2*, and *Twist1* as important regulators of soft palate development (Sugii et al. 2017, Liu et al. 2008, Han et al. 2021). Wnt signaling is of particular interest for further investigation since it is a known mediator of TGF-β signaling during soft palate development (Iwata et al. 2014) but its mechanisms of action in this context have not yet been elucidated. Furthermore, comprehensive screening of expression patterns related to signaling pathways potentially involved in soft palate development led us to the finding that Wnt signaling is predominantly expressed in cells of CNC origin closely surrounding the myogenic soft palatal cells, making it a strong candidate for mediating the communication between CNC-derived mesenchymal and myogenic cells (Janeckova et al. 2019). In general, Wnt signaling is crucial for craniofacial development in humans and mice, and disruptions to Wnt signaling cause severe craniofacial defects, including cleft palate (Brault et al. 2001, Huelsken et al. 2000, Chen et al. 2009, He et al. 2011, Reynolds et al. 2019). Independently, Wnt signaling has also been shown to play a role during various phases of myogenesis (Suzuki et al. 2018, Zhong et al. 2015).

In the present study, using the soft palate as a model, we highlight how canonical Wnt signaling acts upstream of ciliogenesis. Primary cilia have been identified as essential mediators of the complex tissue-tissue interactions accompanying soft palate development. We show that persistence of the primary cilia, caused by their dysfunctional disassembly in the CNC-derived mesenchymal cells due to a lack of canonical Wnt signaling, prevents cell cycle progression. This reduces the number of proliferating CNC-derived mesenchymal cells, influences their differentiation status, and ultimately results in underdevelopment of the soft palatal muscles. Our study emphasizes the connection between Wnt signaling and ciliogenesis in regulating tissue-tissue interactions to control organogenesis.

## Results

### Wnt signaling-dependent tissue-tissue interactions are essential for soft palate development

The complete restoration of soft palatal functions in cleft patients is a demanding task for surgeons since it involves extensive repair of musculature (Monroy et al. 2015). Affected individuals may display misorientation, atrophy, and fibrosis of the muscles along with reduced numbers of muscle cells (Von den Hoff et al. 2018, Monroy et al. 2015, Li et al. 2019). In order to understand how these defects arise, we screened several signaling pathways that could be involved in soft palate development, such as Wnt, Fgf and Hh (Janeckova et al. 2019). From this analysis, Wnt signaling emerged as a strong candidate for mediating the signaling among the CNC-derived mesenchymal and myogenic cells since it was primarily activated in the CNC-derived mesenchymal cells (Janeckova et al. 2019). Previously, it has been suggested that Wnt signaling is involved in soft palatal muscle development (Iwata et al. 2014); however, the detailed mechanism has not been revealed. We therefore directed our focus to the role of canonical Wnt signaling in this process.

To test the hypothesis that canonical Wnt signaling is functionally required for regulating tissue-tissue interactions in controlling soft palate development, we deleted *β-catenin* from CNC-derived mesenchymal cells by using the *Osr2-Cre* driver (Chen et al. 2009), which does not influence epithelial or myogenic cells (Fig. S1A-D). To validate the efficient inactivation of β-catenin in CNC-derived mesenchymal cells, we analyzed its expression at the protein level at E13.5 and confirmed its absence from the CNC-derived mesenchymal cells, as well as the persistence of its expression in epithelial cells, in *Osr2-Cre;β-catenin^fl/fl^* mice in comparison to the control (Fig. S1E-F and G-H). Loss of *β-catenin* in palatal CNC-derived mesenchymal cells resulted in defects in the palatine process of the maxilla and palatine bone in *Osr2-Cre;β-catenin^fl/fl^*mice (Fig. S2A and D). Furthermore, soft tissue microCT scans revealed a complete cleft palate (Fig. S2B and E), which was confirmed in histological sections (Fig. S2C and F). The hard palatal shelves of *Osr2-Cre;β-catenin^fl/fl^* mice failed to elevate fully, grow horizontally toward each other, and fuse. These phenotypes resulted in postnatal lethality and signified that canonical Wnt signaling in the CNC-derived mesenchymal cells is indispensable for palatogenesis.

To further investigate the role of Wnt signaling in regulating soft palate development, we analyzed the soft palatal region in *Osr2-Cre;β-catenin^fl/fl^* mice. The levator veli palatini (LVP) is the main subject of our analysis because it is considered the main muscle of the soft palate (Monroy et al. 2015). In comparison to the control, the soft palatal shelves are not developed at E18.5 in the *Osr2-Cre;β-catenin^fl/fl^* mice (Fig. 1A, E and C, G). The LVP was significantly affected, and no myogenic fibers were detected in the *Osr2-Cre;β-catenin^fl/fl^* mice in comparison to the control at E18.5 (Fig. 1B, F, and D, H). In order to discover when the muscle phenotype first appears in these mice, we analyzed embryos at different stages and found that the palatal shelf development and the number of MyoD positive cells at the level of LVP were comparable between controls and mutants at E13.5 (Fig. 1I-J and K-L). The difference in the number of *Myod1* positive cells started to be apparent in *Osr2-Cre;β-catenin^fl/fl^* mice at E14.0 in comparison to the control (Fig. 1M and O) and persisted at E14.5 (Fig. 1N and P). Therefore, we considered E13.5-E14.0 as the onset of the phenotype, and we used mice at these stages for further analyses. In summary, these results indicated that Wnt signaling is required for tissue-tissue interactions between the CNC-derived mesenchymal cells and mesoderm-derived myogenic cells in regulating soft palate development.

**Fig. 1.**
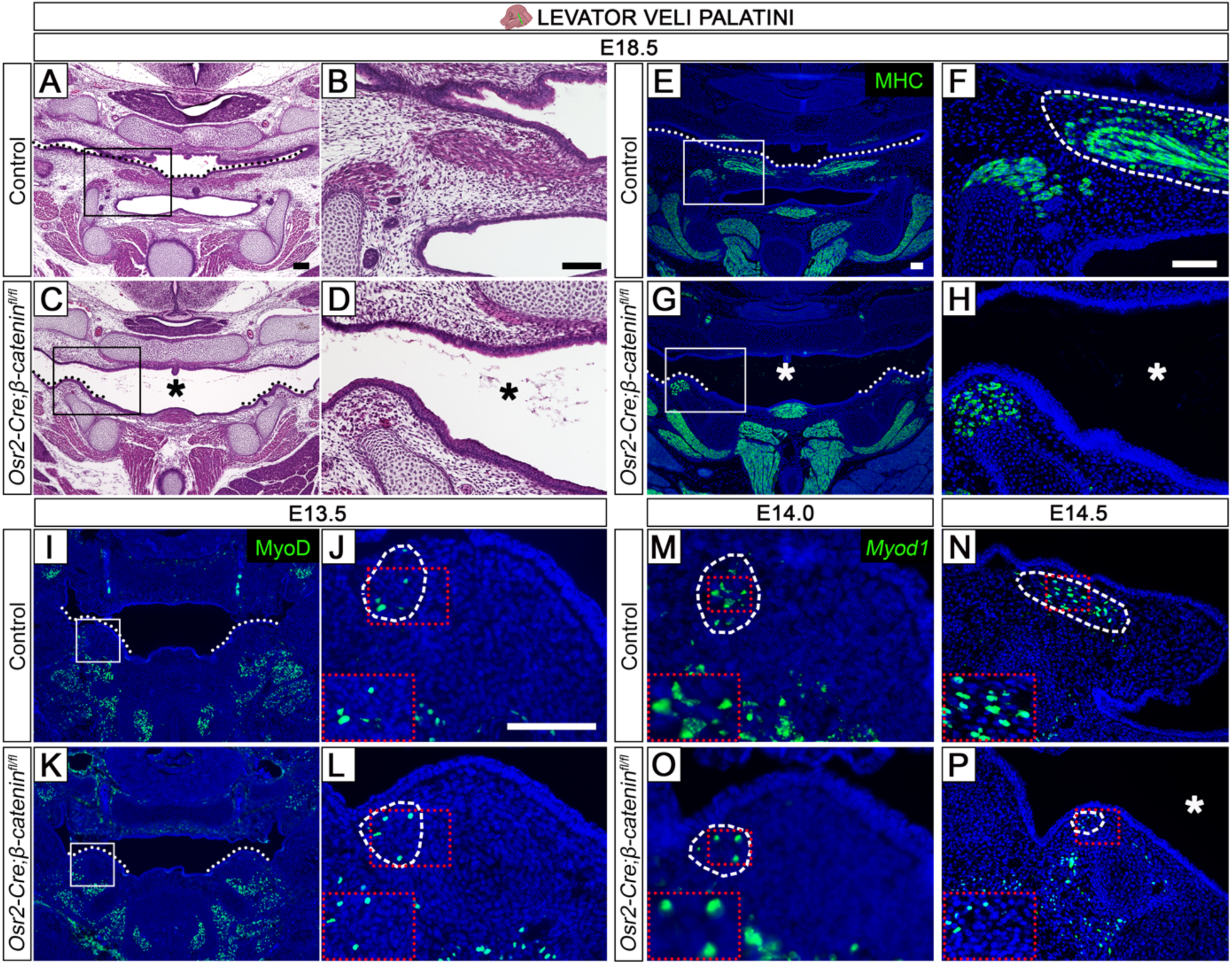
Wnt signaling from CNC-derived mesenchymal cells is essential for palatogenesis and guiding myogenesis through tissue-tissue interactions. (**A-D**) Hematoxylin and eosin staining at the level of the levator veli palatini (LVP) in control (A-B) and *Osr2-Cre;β-catenin^fl/fl^* mice (**C-D**) at E18.5. Black asterisks in C and D identify the cleft palate in *Osr2-Cre;β-catenin^fl/fl^*. (E-H) MHC (green) immunofluorescence in control (E-F) and *Osr2-Cre;β-catenin^fl/fl^* mice (**G-H**) at E18.5. White asterisks in G and H identify the cleft palate in *Osr2-Cre;β-catenin^fl/fl^*. (I-L) MyoD (green) immunofluorescence in control (I-J) and *Osr2-Cre;β-catenin^fl/fl^* mice (K-L) at E13.5. (M-P) *Myod1* (green) *in situ* RNAScope hybridization in control (M-N) and *Osr2-Cre;β-catenin^fl/fl^* mice (**O-P**) at E14.0 (M, O) and E14.5 (N, P). White asterisk in P identifies the cleft palate in *Osr2-Cre;β-catenin^fl/fl^*. Schematic drawing of the mouse head in the top panel depicts the position and angle of sectioning. Black boxes in A and C show approximate locations of higher magnification images in B and D, respectively. White boxes in E, G, I, and K show approximate locations of higher magnification images in F, H, J, M, N, L, O, and P. Red dotted boxes in J, L, and M-P show the position of the insets in their lower left corners. Black dotted lines in A and C indicate palatal shelves. White dotted lines in E, G, I, and K indicate palatal shelves. White dashed lines outline the LVP muscle region stained with MHC, MyoD, or *Myod1* in F, J, L, and M-P. Scale bar in A indicates 200 µm in A, and C; scale bar in B indicates 100 µm in B, and D; scale bar in E indicates 100 µm in E, G, I and K; scale bar in F indicates 100 µm in F, H, N, and P; scale bar in J indicates 100 µm in J, L, M, and O, respectively.

### Wnt signaling is required for the proliferation and cell cycle progression of CNC-derived mesenchymal cells in the soft palate

Balanced cellular activities, such as proliferation and apoptosis, are crucial for mesenchymal cells during soft palate development. In order to determine the cause of the stunted growth of the soft palatal shelves in *Osr2-Cre;β-catenin^fl/fl^* mice, we analyzed cellular activities of CNC-derived mesenchymal cells at E13.5 and E14.0, when the palatal shelves were still comparable between controls and mutants at the level of the LVP, and the palatal shelves consisted predominantly of CNC-derived mesenchymal cells with only a few myogenic cells. Our analysis showed decreased proliferation of soft palatal mesenchymal cells in *Osr2-Cre;β-catenin^fl/fl^* mice at E13.5 (Fig. 2A-B and D-E, I). However, there was no difference in apoptosis between controls and mutants at E13.5 (Fig. 2C and F). To uncover any potential changes in cell cycle progression that could further elucidate the dynamic changes between the control and *Osr2-Cre;β-catenin^fl/fl^*mice, we co-localized the S-phase marker BrdU with M-phase marker pH3. This co-localization identifies cells that progress in the cell cycle from S-to M-phase during the short BrdU labeling. Our analysis revealed fewer BrdU+ pH3+ double-positive cells in the total BrdU+ population in the palatal shelves of *Osr2-Cre;β-catenin^fl/fl^* mice at E14.0 (Fig. 2G-H and J-K, L). Therefore, fewer S-phase cells progressed into M-phase in *Osr2-Cre;β-catenin^fl/fl^*mice in comparison to the control. These results indicated that Wnt signaling has a specific role in maintaining appropriate proliferation and cell cycle progression of CNC-derived mesenchymal cells in the soft palate.

**Fig. 2.**
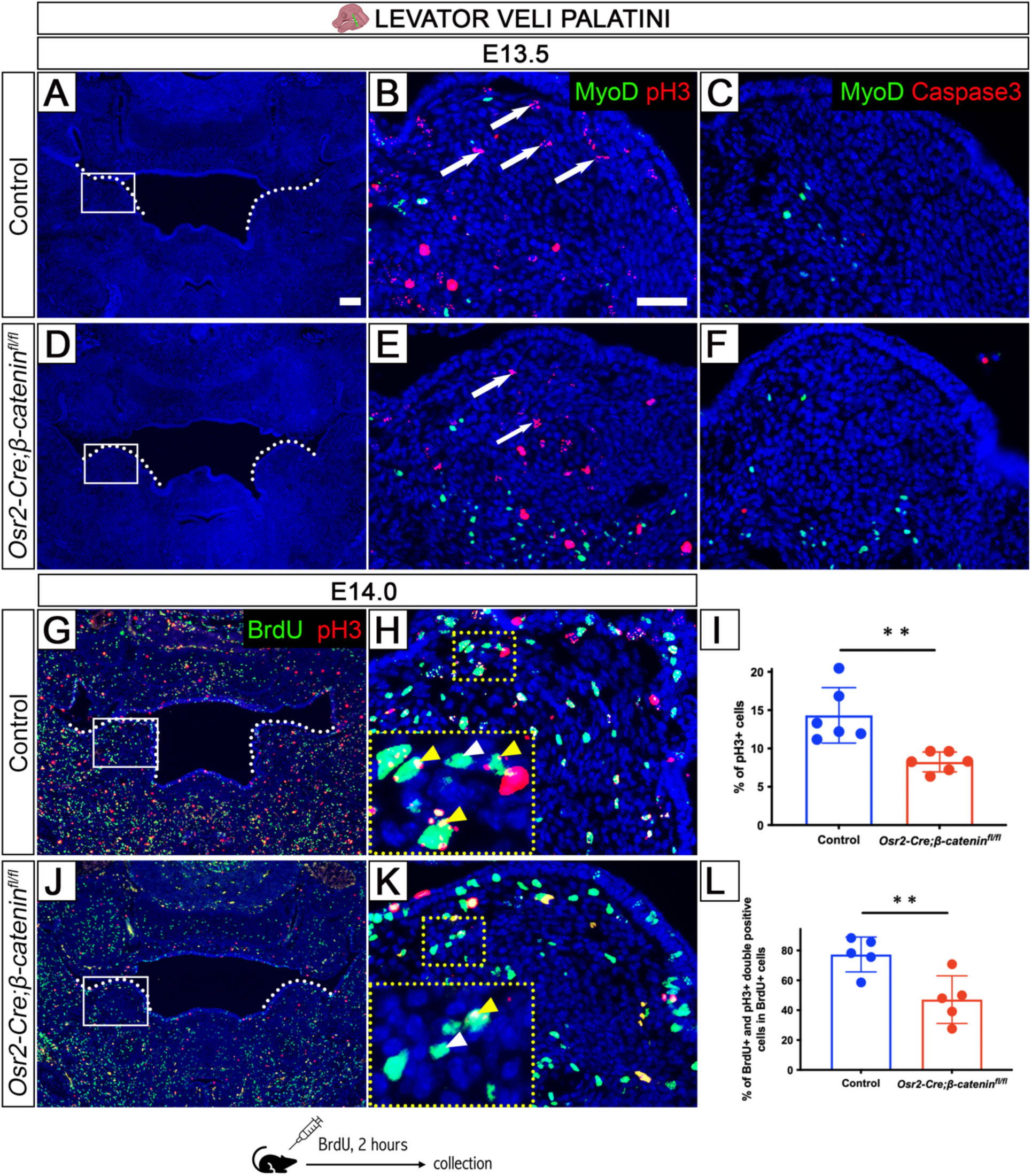
*Osr2-Cre;β-catenin^fl/fl^* mice display decreased proliferation of the mesenchymal cells. (A, D) DAPI staining of control (A) and *Osr2-Cre;β-catenin^fl/fl^* (D) sections at the level of the LVP at E13.5. (B-C, E-F) MyoD (green) and pH3 (red) (B, E), or MyoD (green) and caspase3 (red) immunohistofluorescence staining (C, F) in control (B-C) and *Osr2-Cre;β-catenin^fl/fl^* mice (E-F). (G-H, J-K) BrdU (green) and pH3 (red) immunohistofluorescence staining at E14.0 in control (G-H) and *Osr2-Cre;β-catenin^fl/fl^*mice (J-K). (I) Quantification of the percentage of pH3 positive cells. **p-value = 0.0031. (L) Quantification of percentage of BrdU+ pH3+ double positive cells out of total BrdU+ cells. **p-value = 0.0091. Schematic drawing of the mouse head in the top panel depicts the position and angle of sectioning. Schematic drawing at the bottom indicates injection of pregnant mouse with BrdU followed by collection of embryos 2 hours later. White boxes in A, D, G, and J show approximate location of higher magnification images in B-C, E-F, H, and K, respectively. Yellow dotted boxes in H and K show the position of the insets in their lower left corners. White arrows in B and E point to the positive signal. White arrowheads in H and K point to positive cells. Yellow arrowheads in H and K point to double positive cells. Scale bar in A indicates 100 µm in A, D, G, and J; scale bar in B indicates 50 µm in B-C, E-F, H, and K.

### Absence of Wnt signaling in mesenchymal cells increases ciliogenesis and decreases myogenesis in the soft palatal region

To uncover how canonical Wnt signaling regulates proliferation in the soft palatal mesenchyme, we performed RNA sequencing (RNAseq) analysis of *Osr2-Cre;β-catenin^fl/fl^* and control mice at E14.0 (Fig. 3A-C) to identify changes in the transcriptome and downstream targets with significantly changed expression in the absence of Wnt signaling in soft palatal mesenchymal cells. We identified significant changes in myogenesis and ciliogenesis in the soft palate of *Osr2-Cre;β-catenin^fl/fl^* samples compared with controls (Fig. 3D). The decrease in soft palatal myogenesis in *Osr2-Cre;β-catenin^fl/fl^* mice indicated that Wnt signaling in CNC-derived mesenchymal cells is necessary for myogenic cells through tissue-tissue interactions, consistent with the defects observed in histological analysis of the myogenic cells (Fig. 1A-H). The most striking change was increased ciliogenesis (e.g.: cilium, GO:0005929; axoneme, GO:0005930), which contrasted with the decreased myogenesis and compromised muscle cell development seen in these *β-catenin* mutant mice. Primary cilia are complex and unique organelles that play an important role in organogenesis and function as cellular antennas (Wheway et al. 2018). The potential involvement of primary cilia in soft palate development has not been explored. To comprehensively validate the ciliary phenotype, we analyzed ciliary structure, intraflagellar transport and posttranslational modifications of cilia (Fig. S3A).

**Fig. 3.**
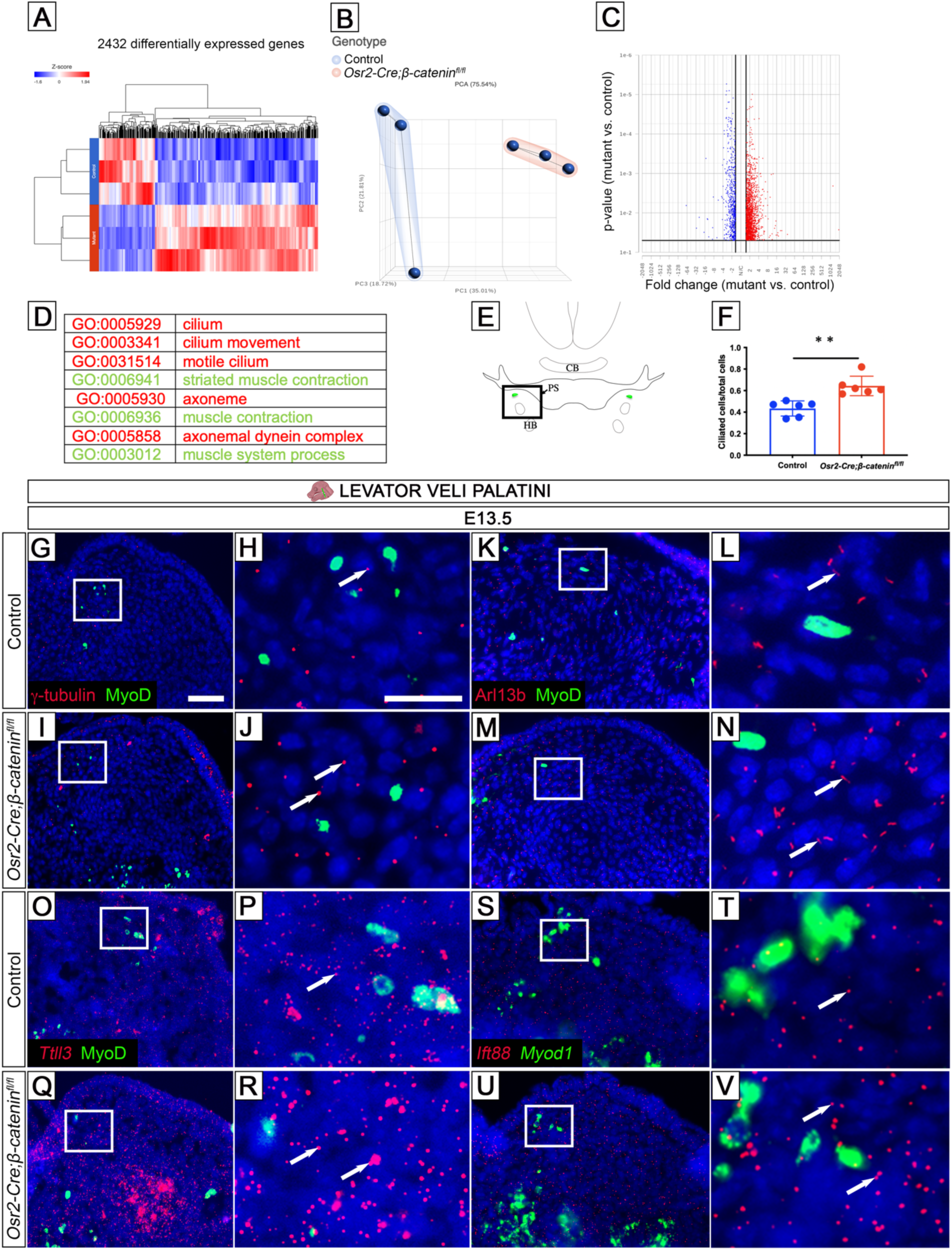
Loss of Wnt signaling in mesenchymal cells leads to an increase in the number of cilia. (A-C) RNAseq analysis of soft palatal samples from control (blue) and *Osr2-Cre;β-catenin^fl/fl^* mice (red) collected at E14.0 depicted with heat map (A), PCA plot (B), and volcano plot (C). RNAseq analysis was performed under the following conditions: p-value ≤ 0.05, fold change ≤-1.5 or ≥ 1.5. A total of 2122 differentially expressed genes were identified, out of which 1810 were upregulated and 622 downregulated. (D) The most changed GO terms from the Partek Flow analysis point to the changes in muscle (green) and cilium development (red). (E) Schematic drawing at the level of the levator veli palatini level (LVP). (F) γ-tubulin quantification indicates the percentage of ciliated cells. **p-value = 0.0012. (G-J) γ-tubulin (red) and MyoD (green) immunohistofluorescence staining at E13.5 in control (G-H) and *Osr2-Cre;β-catenin^fl/fl^* mice (I-J). (K-N) Arl13b (red) and MyoD (green) immunohistofluorescence staining at E13.5 in control (K-L) and *Osr2-Cre;β-catenin^fl/fl^* mice (M-N). (O-R) *Ttll3* (red) *in situ* RNAScope hybridization and MyoD (green) immunohistofluorescence staining in control (O-P) and *Osr2-Cre;β-catenin^fl/fl^* mice (Q-R). (S-V) *Ift88* (red) and *Myod1* (green) *in situ* RNAScope hybridization in control (S-T) and *Osr2-Cre;β-catenin^fl/fl^* mice (U-V). Schematic drawing of the mouse head in the top panel depicts the position and angle of sectioning. Black box in E shows approximate location of images in G, I, K, M, O, Q, S, and U. White boxes in G, I, K, M, O, Q, S, and U are showing a higher magnification in H, J, L, N, P, R, T, and V, respectively. White arrows in H, J, L, N, P, R, T, and V point to the positive signal. Scale bar in G indicates 50 µm in G, I, K, M, O, Q, S, and U; scale bar in H indicates 20 µm in H, J, L, N, P, R, T, and V.

To analyze the ciliary structure in *Osr2-Cre;β-catenin^fl/fl^* mice, we used γ-tubulin (a marker of the ciliary basal body), and Arl13b (a marker of the ciliary axoneme). We identified the increased expression of γ-tubulin and Arl13b in the palatal mesenchyme of *Osr2-Cre;β-catenin^fl/fl^*mice in comparison to the control through immunofluorescence staining and quantification of the ratio of ciliated cells to total cells (Fig. 3F, G-H, K-L and I-J, M-N). Since cilia are known to be dynamically regulated during cell cycle progression (Kasahara and Inagaki 2021) and our previous analysis showed that mesenchymal cells are not progressing to M-phase in the palatal mesenchyme of *Osr2-Cre;β-catenin^fl/fl^* mice, we analyzed the RNAseq data with a focus on genes that are essential for ciliary stability and homeostasis (the balance between assembly and disassembly). One of the genes we identified is *Ttll3*, which encodes a glycylase that serves as a post-translational modifier of the ciliary axoneme (Gadadhar et al. 2017). The increased expression of *Ttll3* was confirmed through RNAScope *in situ* staining of the palatal mesenchyme (Fig. 3O-P and Q-R). Another posttranslational modifier is *Ttll6,* which also showed increased expression in the palatal mesenchyme of *Osr2-Cre;β-catenin^fl/fl^* mice (Fig. S3B-C and D-E). Furthermore, we discovered that the expression of *Ift88,* which is involved in ciliary intraflagellar transport, increased in the palatal mesenchyme of *Osr2-Cre;β-catenin^fl/fl^* mice (Fig. 3S-T and U-V). Increased expression of cilium-related genes, especially that of *Ttll3*, enhances the stability of cilia by preventing their disassembly, which explains the increased number of ciliated cells seen in the palatal mesenchyme of *Osr2-Cre;β-catenin^fl/fl^* mice. In summary, these experiments revealed that, in the absence of canonical Wnt signaling in mesenchymal cells, myogenesis and ciliogenesis are adversely affected in *Osr2-Cre;β-catenin^fl/fl^*mice.

### *Ttll3* expression is directly regulated by Tcf7l2/β-catenin complex

Since *Ttll3* has been identified as the key factor responsible for dysfunctional ciliary disassembly in *Osr2-Cre;β-catenin^fl/fl^* mice, we further focused on this gene. To uncover how the expression of *Ttll3* is controlled by canonical Wnt signaling, we searched its promoter region for potential binding sites of the canonical Wnt-related transcription factors Tcf7, Tcf7l1, Tcf7l2, and Lef1, all of which contain a β-catenin binding domain (Cadigan and Waterman 2012). Binding of β-catenin to these transcription factors in the nucleus further enables activation or repression of Wnt-responsive genes. Bioinformatic analysis using transcription binding profiles from the UCSC/JASPAR database showed that Tcf7 and Tcf7l2 can potentially bind to the *Ttll3* promoter region (Fig. 4A). This indicates that *Ttll3* may act downstream of canonical Wnt signaling to control the ciliary disassembly on mesenchymal cells and ultimately influence soft palate development.

**Fig. 4.**
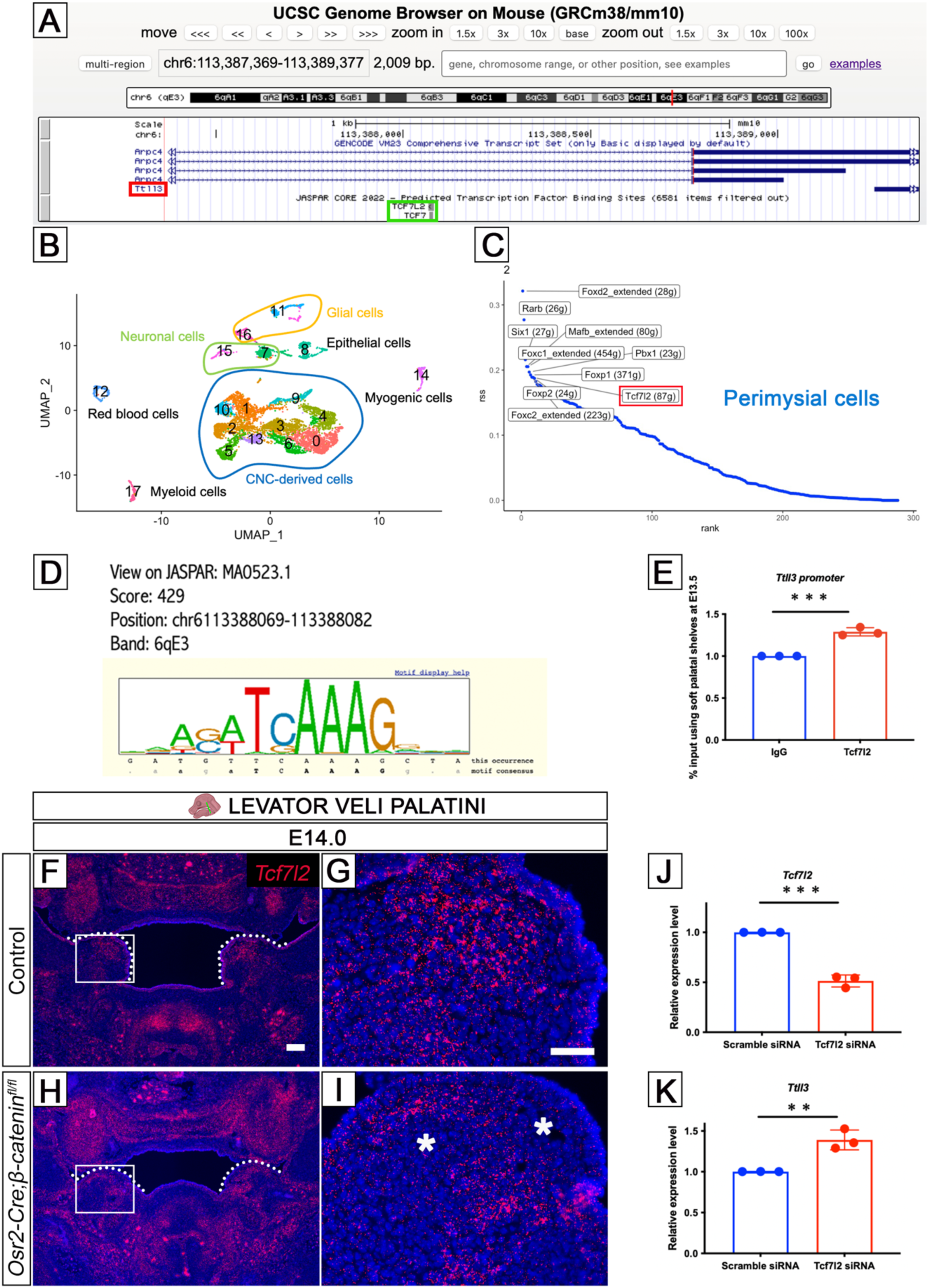
Wnt signaling directly regulates ciliary genes. (A) Predicted position of Tcf7l2 and Tcf7 binding to the promoter region of *Ttll3* from UCSC Genome Browser on Mouse. The red box marks *Ttll3* gene and green box highlights Tcf7l2 and Tcf7. (B) UMAP plot of soft palatal cell populations identified by Seurat from E13.5 soft palatal scRNAseq analysis. 0, midline mesenchymal cells; 1, CNC-derived progenitor cells; 2, perimysial cells; 3, osteogenic cells; 4, perimysial cells; 5, chondrogenic cells; 6, mitotic cells; 7, neuronal cells; 8, epithelial cells; 9, perimysial cells; 10, midline mesenchymal cells; 11, glial cells; 12, red blood cells; 13, chondrogenic cells; 14, myogenic cells; 15, neuronal cells; 16, glial cells; 17, myeloid cells. (C) SCENIC analysis of E13.5 soft palatal scRNAseq data shows Tcf7l2 as an important regulon in perimysial cells. The red box highlights Tcf7l2. (D) UCSC binding prediction of Tcf7l2 binding motif to the promoter of *Ttll3*. (E) Cut and Run assay for predicted binding of Tcf7l2 to *Ttll3* promoter region in the soft palatal tissue at E13.5. ***p-value = 0.0005. (F-I) *Tcf7l2* (red) *in situ* RNAScope hybridization at E14.0 in control (F-G) and *Osr2-Cre;β-catenin^fl/fl^*mice (H-I). (J-K) qPCR analysis. Efficiency of *Tcf7l2* siRNA. ****p-value = 0.0001 (J). Increased expression of *Ttll3* after *Tcf7l2* siRNA treatment. **p-value = 0.0053 (K). Schematic drawing of the mouse head in the top panel depicts the position and angle of sectioning. White boxes in F and H show approximate location of higher magnification images in G and I, respectively. White dotted lines indicate palatal shelves in F and H. Asterisks in I show a lack of positive signal. Scale bar in F indicates 100 µm in F and H; scale bar in G indicates 50 µm in G and I.

To determine if Tcf7 or Tcf7l2 would be better candidate for further analysis and to identify the main genetic regulatory networks in early soft palate development, we analyzed our scRNAseq data from E13.5 soft palatal shelves (Fig. 4B) using the R package SCENIC (Aibar et al. 2017). We revealed that the Tcf7l2 regulon is present in perimysial cells (Fig. 4C), a population which closely interacts with myogenic progenitor cells during the early development of the soft palate and is essential for further myogenic development (Han et al. 2021). These data further support the importance of canonical Wnt signaling in early soft palate development and point to Tcf7l2 being an important transcription factor during this process. We identified a motif (chr6113388069-113388082) that enables binding of Tcf7l2 to the *Ttll3* promoter region (Fig. 4D). Furthermore, we confirmed the direct binding of Tcf7l2 to the predicted region at the *Ttll3* promoter site using a Cut and Run assay (Fig. 4E), suggesting that *Ttll3* is a direct target of canonical Wnt signaling in the soft palate.

Next, we examined the expression of *Tcf7l2* in the soft palatal region *in vivo*. *Tcf7l2* was expressed primarily in the CNC-derived palatal mesenchymal cells in the control and this expression was decreased in *Osr2-Cre;β-catenin^fl/fl^*mice (Fig. 4F-G and H-I). To confirm whether Tcf7l2 serves as the canonical Wnt signaling-related transcription factor responsible for changes in *Ttll3* expression, we performed *in vitro* analysis on cells isolated from the E13.5 control soft palates. Following efficient siRNA-mediated knockdown of *Tcf7l2* in primary soft palatal mesenchymal cells (Fig. 4J), the expression of *Ttll3* was significantly increased (Fig. 4K), suggesting that Tcf7l2 acts as a repressor of *Ttll3* expression in soft palatal mesenchymal cells. In summary, Tcf7l2 can functionally regulate the expression of *Ttll3* by directly binding to its promoter and decreased levels of β-catenin*/*Tcf7l2 in CNC-derived mesenchymal cells cause stabilization of cilia and thus prevent the progression of these cells in the cell cycle.

Thus, in normal conditions, the β-catenin*/*Tcf7l2 complex functions as a repressor of the ciliary gene *Ttll3*, which ultimately influences ciliary stability; the increased expression of *Ttll3* in *Osr2-Cre;β-catenin^fl/fl^* mice causes dysfunctional ciliary disassembly. These results reveal the mechanism by which Wnt signaling regulates the precise balance of ciliary assembly/disassembly that is indispensable for soft palate development.

### Ciliary assembly restrains cell cycle progression in soft palatal mesenchymal cells

Our results suggest that the disturbed cell cycle progression of the palatal mesenchymal cells in *Osr2-Cre;β-catenin^fl/fl^* mice is caused by dysfunctional ciliary disassembly, disrupting ciliary homeostasis. Therefore, we hypothesized that restoring the ciliary homeostatic balance could rescue the proliferation defect. To test this hypothesis, we treated soft palatal mesenchymal cells isolated from *Osr2-Cre;β-catenin^fl/fl^* mice with *Ttll3* siRNA (targeting the expression of *Ttll3*, a glycylase important for ciliary stability). In cultures treated with 50 nM *Ttll3* siRNA, we observed that the percentage of ciliated cells was significantly decreased, as shown by the percentage of cells exhibiting staining of the ciliary axoneme with Arl13b (Fig. 5A-E). This treatment also led to an increase in proliferation of the CNC-derived soft palatal cells (Fig. 5A-D, F), suggesting that ciliary assembly can restrain cell cycle progression of these cells. More importantly, *Ttll3* siRNA treatment was able to restore the proliferation of CNC-derived palatal mesenchymal cells isolated from *Osr2-Cre;β-catenin^fl/fl^* mice to normal levels (Fig. 5G-K). Collectively, these results indicate that strict control of ciliary homeostasis is essential for proper proliferation, cell cycle progression, and development of the soft palate.

**Fig. 5.**
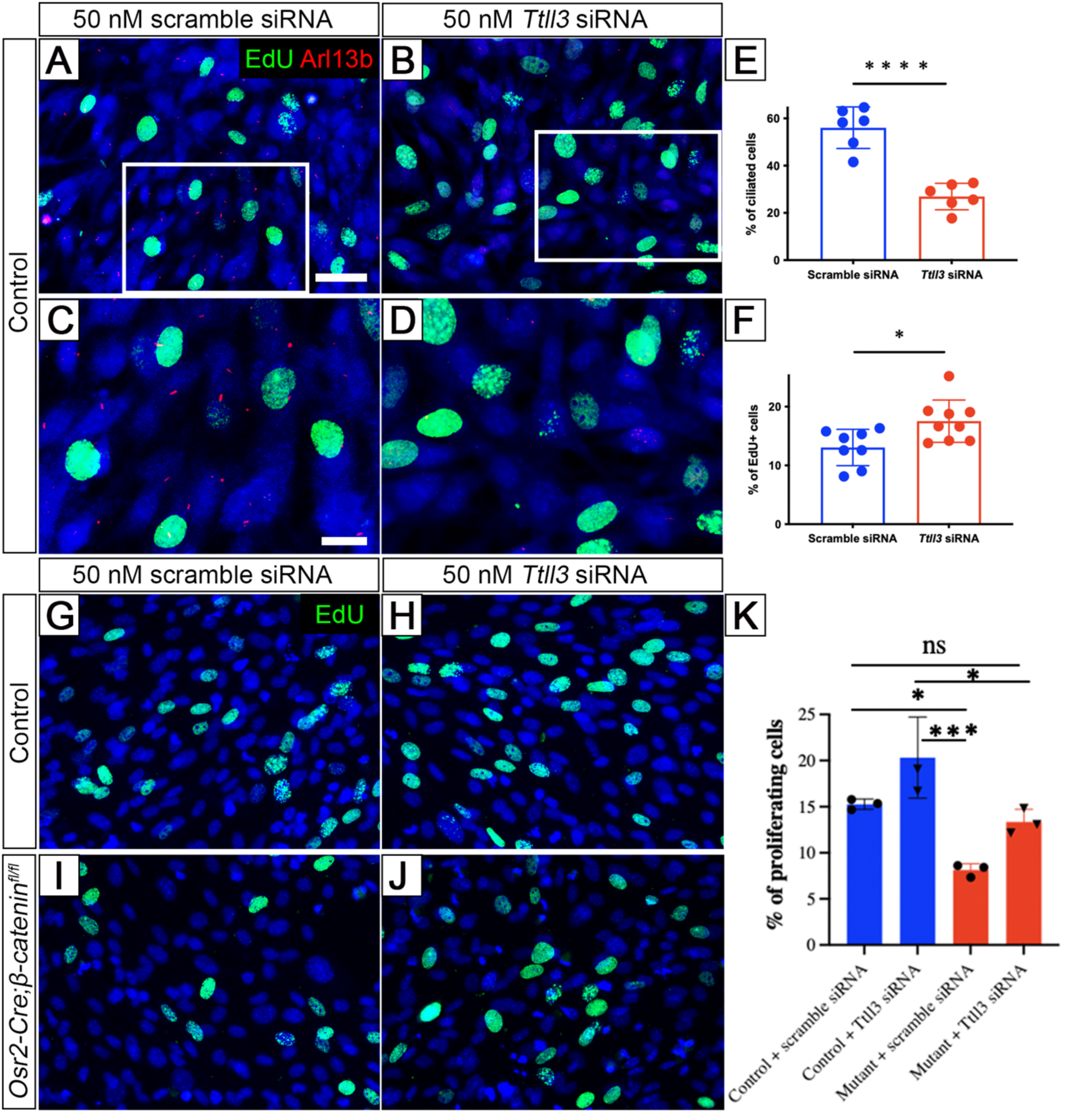
Dysfunctional ciliary disassembly restrains cell cycle progression of soft palatal cells in *Osr2-Cre;β-catenin^fl/fl^* mice. (A-D) EdU (green) and Arl13b (red) immunofluorescence staining on cells isolated from control mice treated with 50 nM scramble siRNA (A, C) or 50 nm *Ttll3* siRNA (B, D). (E) Quantification of percentage of ciliated cells. ****p-value < 0.0001. (F) Quantification of percentage of EdU+ cells. *p-value = 0.0153. (G-J) EdU (green) immunohistofluorescence staining on cells isolated from control (G-H) and *Osr2-Cre;β-catenin^fl/fl^* mice (I-J) treated with 50 nM scramble siRNA (G, I), or 50 nm *Ttll3* siRNA (H, J). (K) Quantification of percentage of EdU+ cells. All p-values ≤ 0.05; ns = not significant. Scale bar in A indicates 50 µm in A-B, and G-J; scale bar in C indicates 20 µm in C-D.

### Primary cilia regulate the perimysial fate of CNC-derived palatal mesenchymal cells

To discover the mechanism of the communication among CNC-derived mesenchymal and myogenic cells, we analyzed our E13.5 soft palatal scRNAseq data because the altered cell proliferation phenotype in *Osr2-Cre;β-catenin^fl/fl^* mice appears just after this stage. We utilized the R package CellChat to identify the predicted ligand-receptor interactions between the CNC-derived mesenchymal and myogenic cells (Jin et al. 2021). Among the highly ranked predicted outgoing signaling patterns from the perimysial cells (clusters 2, 4, and 9) was Notch signaling (Fig. 6A), especially the Dlk1/Notch3 ligand and receptor combination (Fig. 6B), which represents non-canonical Notch signaling that could be received by the myogenic cells. Supporting this notion, we found that the expression of *Dlk1* was decreased in the palatal mesenchyme of *Osr2-Cre;β-catenin^fl/fl^*mice (Fig. 6C-D and E-F). Since primary cilia function as a signaling hub, we hypothesized that, apart from influencing the cell cycle and its progression, primary cilia could also be responsible for the differentiation status of the perimysial cells that are essential for the development of soft palatal muscles. Furthermore, in our scRNAseq analysis, *Dlk1* has been identified as one of the top three molecules enriched in the perimysial cells and it can function both as a transmembrane and secreted protein, making it a good candidate for mediating the communication among CNC-derived mesenchymal and myogenic cells. After sufficiently decreasing the expression of *Ttll3* using siRNA in the primary soft palatal mesenchymal cells *in vitro* (Fig. 6G), we observed increased expression of *Dlk1* (Fig. 6H). These findings are consistent with our *in vivo* finding that overexpression of *Ttll3* caused a decrease of *Dlk1* expression. Therefore, in the soft palate, primary cilia coordinate both the proliferation and cell cycle progression of the CNC-derived mesenchymal cells as well as the differentiation of perimysial cells.

**Fig. 6.**
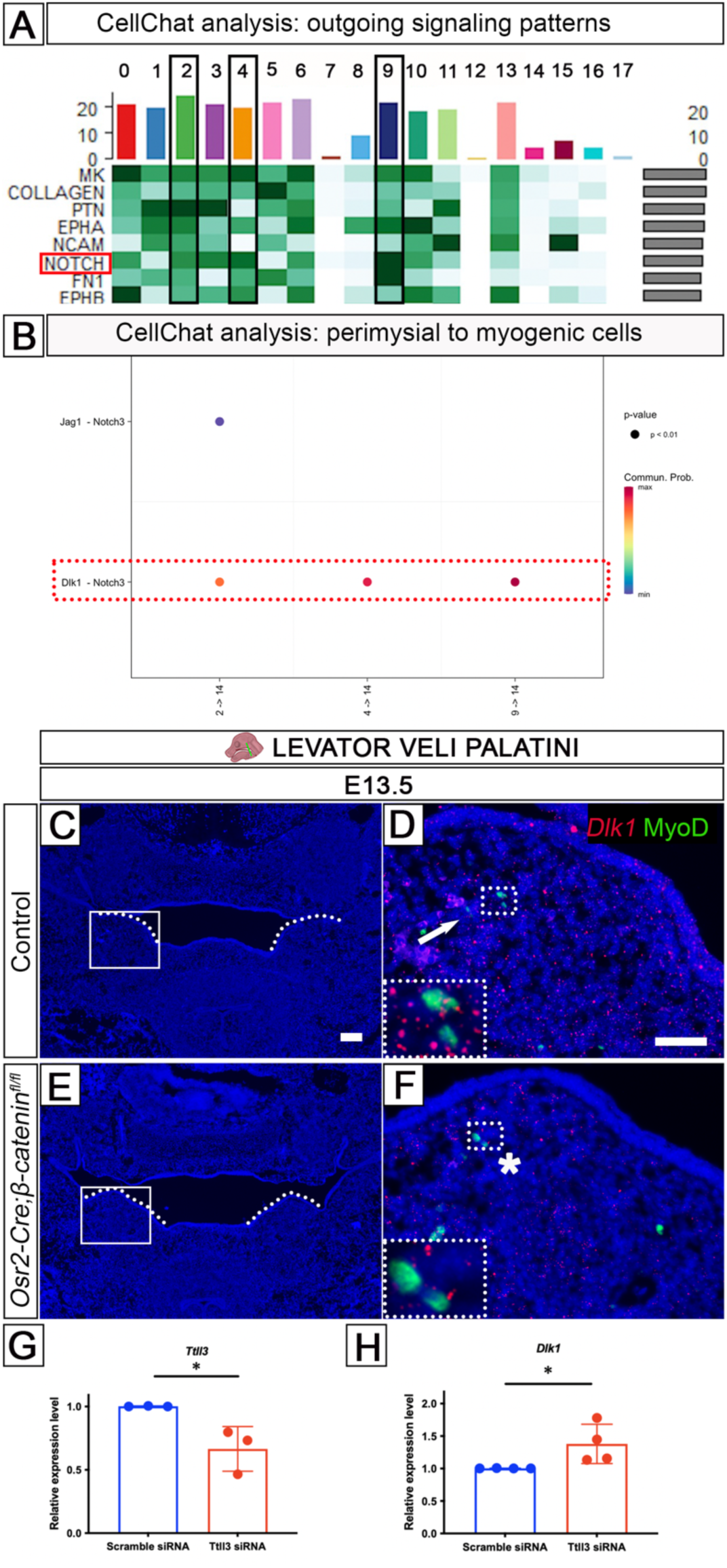
Primary cilia influence the soft palate muscle development via regulating the perimysial fate of CNC-derived palatal mesenchyme cells, which further disrupts soft palate myogenesis. (A) CellChat analysis of E13.5 soft palatal scRNAseq data showing a heatmap visualizing the contribution of the outgoing signaling patterns. 0, midline mesenchymal cells; 1, CNC-derived progenitor cells; 2, perimysial cells; 3, osteogenic cells; 4, perimysial cells; 5, chondrogenic cells; 6, mitotic cells; 7, neuronal cells; 8, epithelial cells; 9, perimysial cells; 10, midline mesenchymal cells; 11, glial cells; 12, red blood cells; 13, chondrogenic cells; 14, myogenic cells; 15, neuronal cells; 16, glial cells; 17, myeloid cells. The red box highlights Notch signaling and black boxes mark perimysial clusters (2, 4, 9). (B) Notch signaling activity is shown in the direction of perimysial to myogenic cells (2→14, 4→14, and 9→14). Red dotted box highlights Dlk1/Notch3 ligand and receptor combination. (C-F) *Dlk1* (red) *in situ* RNAScope hybridization and MyoD (green) immunohistochemistry staining at E13.5 in control (C-D) and *Osr2-Cre;β-catenin^fl/fl^* mice (E-F). (G-H) qPCR analysis. Efficiency of *Ttll3* siRNA. *p-value = 0.0297 (G). Increased expression of *Dlk1* after *Ttll3* siRNA treatment. *p-value = 0.0477 (H). Schematic drawing of the mouse head in the top panel depicts the position and angle of sectioning. White boxes in C and E show approximate location of higher magnification images in D and F, respectively. White dotted boxes in D and F show the position of the insets in their lower left corners. White dotted lines indicate palatal shelves in C and E. White arrow in D points to positive signal. Asterisk in F shows a lack of positive signal. Scale bar in C indicates 100 µm in C and E; scale bar in D indicates 50 µm in D and F.

## Discussion

The soft palate is an essential part of the oropharyngeal complex, fulfilling vital functions including breathing, hearing, swallowing and speech, all of which are affected in cleft patients (Monroy et al. 2015). Cleft soft palate is less commonly studied than cleft hard palate, yet it accompanies the majority of cleft cases, and functional restoration of the palatal soft tissue has a low surgical success rate (Monroy et al. 2015, Von den Hoff et al. 2018). Moreover, it imposes a significant burden on patients, families, and healthcare providers since multidisciplinary treatment may be required into adulthood (Von den Hoff et al. 2018). Clinically, patients with soft palatal clefts exhibit a reduced number of myogenic cells, together with misorientation, lack of fusion, misalignment, atrophy, and fibrosis of the soft palatal muscles (Li et al. 2019, Monroy et al. 2015). The detailed molecular mechanisms involved in regulating soft palate development still need to be elucidated, including the reciprocal interactions among the CNC-derived mesenchymal cells, mesoderm-derived myogenic cells, ectoderm-derived epithelial cells, and mediators of these communication channels, such as the secreted factors mediating the message or special organelles that may be involved (Li et al. 2019). A better understanding of these signaling mechanisms may contribute to the development of improved, biologically informed soft palatal muscle restoration strategies.

After screening for the expression patterns of several signaling pathways, we uncovered that Wnt signaling is activated primarily in the CNC-derived mesenchymal cells surrounding the myogenic cells during soft palatal muscle development. While mutations in Wnt signaling genes are well-described in orofacial clefting in humans as well as mice, and this pathway is independently known to play a role in myogenesis; here, we focused on its function in the soft palate (Reynolds et al. 2019, Suzuki et al. 2018). Studying Wnt signaling in murine palatogenesis is otherwise very challenging due to embryonic lethality and failure of craniofacial development after conditional deletion of the *β-catenin* gene using the more broadly expressed *Wnt1-Cre* (Brault et al. 2001). In the current study, narrowly targeted conditional knockout of canonical Wnt signaling only from the CNC-derived mesenchymal cells using *Osr2-Cre* created an opportunity to study the role of Wnt signaling in regulating tissue-tissue interactions during soft palate development.

We discovered that canonical Wnt signaling is essential for proper soft palatal muscle development, and in particular, its regulation of ciliogenesis is indispensable for the proliferation and cell cycle progression of CNC-derived mesenchymal cells and their differentiation. These findings are in agreement with previously published data establishing that cilia need to disassemble before the cells enter the M-phase of the cell cycle; otherwise, they can function as a cell cycle brake (Liu et al. 2021). Indeed, in *Osr2-Cre;β-catenin^fl/fl^* mice the mesenchymal cells do not progress into M-phase and exhibit increased expression of *Ttll3* which encodes a glycylase that is required for ciliary stability (Gadadhar et al. 2017). Restoring the ciliary homeostasis by knocking down the expression of *Ttll3* rescues these proliferation defects. Canonical Wnt signaling therefore influences ciliary posttranslational modifications and disassembly, and ultimately the cell cycle progression of the CNC-derived mesenchymal cells, in the model of the soft palate.

Since canonical Wnt signaling is a complex pathway, it is important to determine which of its mediators is the furthest downstream, such that it can bind with β-catenin to directly regulate the expression of ciliary genes. Transcription factors Tcf7, Tcf7l1, Tcf7l2, and Lef1 each contain a β-catenin binding domain and this binding determines the specificity of canonical Wnt signaling in activating or repressing canonical Wnt-responsive genes in various tissues and organs (Cadigan and Waterman 2012). We established that β-catenin/Tcf7l2 complex can regulate genes related to ciliogenesis, which plays an important role in regulating the integrity of the soft palatal shelves and muscle development. Moreover, Tcf7l2 has been identified as an important regulon playing a role in perimysial cells, further supporting the importance of canonical Wnt signaling in early soft palate development. This data is consistent with the previously reported role of Tcf7l2-mediated canonical Wnt signaling in early embryonic limb muscle development (Kardon et al. 2003). Also, Tgf-β-driven reduction of Tcf7l2 occurs in the mesenchymal stromal cells but not in myoblasts, also supporting the role of Tcf7l2 in cells surrounding the muscle cells (Contreras et al. 2020).

Intriguingly, the relationship between Wnt signaling and ciliogenesis is unclear. To date, limited studies have reported Wnt signaling being upstream of ciliogenesis. In zebrafish it was demonstrated on the basis of Kupffer’s vesicle motile cilia (Caron et al. 2012) and *in vitro* experiments on hTERT RPE cells (human telomerase reverse transcriptase-immortalized retinal pigment epithelial cell line) that Wnt3a stimulation promotes primary ciliogenesis (Kyun et al. 2020). The effects of cilia on Wnt signaling in the reverse direction are uncertain. After manipulating cilium-related genes, Wnt signaling may be restricted (Corbit et al. 2008) or unchanged (Ocbina et al. 2009), and it has also been proposed that cilia serve as a switch between canonical and non-canonical Wnt signaling (Simons et al. 2005). Our study sheds light on the intricate relations between Wnt signaling and ciliogenesis. We demonstrate the direct involvement of Wnt signaling pathway in the regulation of ciliary genes during soft palate development.

Furthermore, we have discovered a previously unappreciated role of primary cilia in regulating soft palate development. Several studies have reported on craniofacial phenotypes, including cleft palate, associated with ciliopathies, especially Joubert syndrome, Meckel-Gruber syndrome, Oro-facial-digital syndrome, and Ellis-van Creveld syndrome (Brugmann et al. 2010b, Schock and Brugmann 2017). Arl13, which marks the ciliary axoneme, showed an increased number of cilia in CNC-derived soft palatal mesenchymal cells in *Osr2-Cre;β-catenin^fl/fl^*mice and is linked to Joubert syndrome manifesting with cleft palate (Cantagrel et al. 2008, Poretti et al. 2012, Caspary et al. 2007). So far, primary cilia have been shown to play a role in hard palate (*Ift88*, *Kif3a* and *Tmem107*) and skeletal muscle development (Brugmann et al. 2010a, Fu et al. 2014, Tian et al. 2017), but whether they are involved in soft palate development has not been reported. Our study highlights primary cilia as key mediators of soft palatal muscle development.

Cilia have also been suggested as key regulators for cell-cell interactions (Wang and Barr 2018). Our further analysis has shown that there is a strict interdependence of individual signals and collective corresponding cellular responses, with primary cilia serving as the integration hub. Previously, cilia have been identified as a key organelle keeping the balance between proliferation and differentiation, for example in kidney epithelial cells; as well as fulfilling this function through cilia-mediated paracrine signaling (Irigoin and Badano 2011, Boletta et al. 2000). In our model, primary cilia are an important connecting factor influencing both the proliferation and differentiation of CNC-derived mesenchymal cells to establish and maintain the relations with myogenic cells through subsequent stages of morphogenesis. In this study, Notch signaling has been identified as one of the top enriched signaling pathways involved in interactions between perimysial-derived signaling molecules communicating with the myogenic cells at the E13.5 stage of soft palate development. Moreover, *Dlk1* has been identified as one of the top three molecules enriched in perimysial cells. We discovered decreased expression of *Dlk1*, suggesting changes in differentiation of perimysial cells, ultimately leading to defective palatal muscle development. This data agrees with previously published work supporting the role of *Dlk1* in skeletal muscle development and regeneration (Waddell et al. 2010, Shin et al. 2014). This understanding of the interplay among canonical Wnt signaling, cilia and, *Dlk1* may be the key to overcoming clinical challenges in the functional restoration of the soft palatal muscles.

Overall, our study points to the complexity and combination of signaling pathways, cellular processes, and their interaction with organelles such as cilia coming together to orchestrate embryonic development of the soft palatal region. Canonical Wnt signaling influences the proliferation and ciliary homeostasis of soft palatal mesenchymal cells, enabling their differentiation and ultimately promoting proper muscle development. Our study describes primary cilia as the missing mechanistic link that relays diverse signals coming from CNC-derived mesenchymal cells to myogenic cells to enable the congruency of the whole system, including the interactions that initiate sustained differentiation of the CNC-derived mesenchymal cells and their functional diversity. These results represent important findings that ultimately will lead to more effective functional muscle restoration for patients with clefting.

## Materials & Methods

### Animals and embryo collection

All animal handling was performed according to federal regulations with approval from the Institutional Animal Care and Use Committee (IACUC) at the University of Southern California (protocol #9320). C57BL/6J mice (Jackson Laboratory, #000664) were used for this study. *Osr2-CreKI* mice (Chen et al. 2009) (Rulang Jiang, Cincinnati Children’s Hospital) were crossed with *β-catenin^fl/fl^* (Brault et al. 2001) (Jackson Laboratory, #004152) and *tdTomato* conditional reporter mice (Jackson laboratory, #007905) to generate *Osr2-Cre;β-catenin^fl/+^* males or *Osr2-Cre;tdTomato^fl/+^* embryos. The *Osr2-Cre;β-catenin^fl/+^* males were further crossed with *β-catenin^fl/fl^*females to generate *Osr2-Cre;β-catenin^fl/fl^* embryos. As control samples, either *β-catenin^fl/+^*or *β*-*catenin^fl/fl^* were used. Pregnant females were euthanized by carbon dioxide overdose followed by cervical dislocation. Embryos (regardless of their sex) were collected at various developmental stages (embryonic day [E]13.5, E14.0, E14.5, E18.5, and postnatal day [P]0) and fixed overnight in 10% neutral buffered formalin solution (Sigma, HT501128). E18.5, and P0 samples were decalcified in ethylenediaminetetraacetic acid (EDTA, pH 7.1–7.3, Alfa Aesar, A15161) for 1 week and processed for cryosectioning or paraffin embedding and sectioning (8 μm).

### MicroCT analysis

MicroCT scans of P0 *β-catenin^fl/fl^* and *Osr2-Cre;β-catenin^fl/fl^* mice were obtained from a SCANCO mCT50 scanner at the University of Southern California Molecular Imaging Center with the X-ray source at 70 kVp and 114 mA, and all the data were collected at a resolution of 10 mM.

### Histological staining and immunohistofluorescence

Paraffin sections were used for hematoxylin and eosin staining. The histological protocol is described in detail in our previous publication (Janeckova et al. 2019). For immunohistofluorescence, cryo-sections were air-dried at room temperature and then at 37°C for 15 minutes each. Distilled water was used for removing OCT (5-minute wash) and slides were treated with 3% hydrogen peroxide solution (10 minutes), citrate-based antigen unmasking solution (Vector, H-3300-250) for 15 minutes at 99°C, blocking reagent (Thermo Fisher Scientific, B40922) for 1 hour at room temperature, and primary antibody overnight, followed by washes in PBST. Incubation with secondary antibody was performed for 2 hours at room temperature with three subsequent washes in PBST and final counterstaining with DAPI (Thermo Fisher Scientific, D1306) for 5 minutes. Primary antibodies used in this study were as follows: rabbit monoclonal active β-catenin (Cell Signaling, 19807S, 1:100 with TSA), mouse monoclonal, myosin heavy chain, MHC (DSHB, MF20, 1:10), rabbit polyclonal anti-phospho-histone H3, pH3 (Sigma Aldrich, 06-570, 1:100), rabbit monoclonal cleaved caspase-3 (Cell Signaling, 9664S, 1:100), rabbit monoclonal MyoD (Abcam, ab133627, 1:200 with TSA), rabbit polyclonal, γ-tubulin (Sigma Aldrich, T5192, 1:100), rabbit polyclonal Arl13b (Proteintech, 17711-1-AP, 1:100), rabbit polyclonal red fluorescein protein, RFP (Rockland, 600-401-379, 1:500 with TSA), rat monoclonal BrdU (Abcam, ab6326, 1:100). The following secondary antibodies were used in this study: Alexa Fluor 488 anti-mouse (Thermo Fisher Scientific, A11001, 1:200), Alexa Fluor 568 anti-rabbit, (Thermo Fisher Scientific, A11011, 1:200), anti-rabbit HRP (Vector Laboratories, PI-1000, 1:200). For active β-catenin, MyoD and RFP immunostaining, TSA Plus FITC or Cy3 (Akoya Bioscience, NEL7744001KT or NEL771B001KT) was applied for 3 minutes before DAPI counterstaining.

### RNAScope

Cryo-sections were air dried at room temperature and at 37°C for 15 minutes each. Distilled water was used for removing the OCT (5-minute wash) and slides were treated with a pre-heated target retrieval reagent (Advanced Cell Diagnostics, ACD, 322000) for 15 minutes at 99°C. Standard RNAScope reagents were used according to the manufacturer’s instructions using RNAScope Multiplex Fluorescent Detection Kit v2 (Advanced Cell Diagnostics, 323110). The following probes from Advanced Cell Diagnostics were used in this study: Mm-*Myod1* (316081), Mm-*Ttll3* (586791), Mm-*Ttll6* (570931), Mm-*Ift88* (420211), Mm-*Tcf7l2* (466901), and Mm-*Dlk1* (405971).

### BrdU and EdU incorporation and staining

A short 2-hour pulse injection of BrdU (500 μg per 10 g body weight) was administered intraperitoneally at E14.0 to *β-catenin^fl/fl^* and *Osr2-Cre;β-catenin^fl/fl^* mice. EdU (10 mg/1 ml PBS) was used on cell cultures for 2 hours before the fixation of the cells overnight in 10% neutral buffered formalin solution (Sigma, HT501128). EdU was visualized using a Click-iT Plus EdU Cell Proliferation Kit (Thermo Fisher Scientific, C10637) followed by DAPI staining (Thermo Fisher Scientific, D1306, 5 minutes).

### RNA sequencing and qPCR Analysis

The posterior part of the secondary palate (last third of the secondary palate, i.e., the region of the soft palate) was dissected into RNALater (Thermofisher Scientific, AM7020). Samples were stored at -80°C, and when sufficient control and mutant samples were obtained (minimum of N = 3 per genotype), RNeasy Plus Micro Kit (Qiagen, 74034) was used to isolate RNA. UCLA Technology Center for Genomics and Bioinformatics performed determination of the RNA quality and subsequent cDNA library preparation and sequencing (single-end reads with 75 cycles, Illumina Nextseq 500 platform). Differential expression was calculated by selecting transcripts with fold change ≤ -1.5 or ≥ 1.5 and a significance level of p <0.05 using PartekFlow. The RNA sequencing data used in this study have been deposited in NCBI’s Gene Expression Omnibus (Edgar et al. 2002) and are accessible through GEO series accession number GSE208619. The soft palatal tissue was also used for qPCR; after RNA isolation, cDNA was transcribed using iScript™ cDNA Synthesis Kit (Bio-Rad, 1708891). qPCR reaction was run using SsoFast™ EvaGreen^®^ Supermix (Bio-Rad, 172-5202) on a Bio-Rad CFX96 Real-Time System. Values were normalized to *Gapdh* using the 2^−ΔΔCt^ method. The primer sequences used in this study were obtained from PrimerBank (Wang et al. 2012) and produced by IDT (Integrated DNA Technologies) as follows: *Gapdh* (forward primer 5’-AGGTCGGTGTGAACGGATTTG-3’, reverse primer 5’-TGTAGACCATGTAGTTGAGGTCA-3’), *Tcf7l2* (forward primer 5’-GCACACATCGTTCAGAGCC-3’, reverse primer 5’-GGGTGTAGAAGTGCGGACA-3’), *Ttll3v2* (forward primer 5’-ATGGGCCGACTCAGAAACG-3’, reverse primer 5’-CGCAAGAGACACCGGATCA-3’), *Dlk1* (forward primer 5’-CCCAGGTGAGCTTCGAGTG-3’, reverse primer 5’-GGAGAGGGGTACTCTTGTTGAG-3’).

### Soft palatal primary cell culture

The soft palate from E13.5 embryos was dissected and digested using Multi Tissue Dissociate Kit 3 (Miltenyi Biotec, 130-110-204) at 37°C (600 rpm for 15 minutes) in a Thermomixer (Thermo Fisher Scientific, 2231000269). After adding the medium to stop the digestion process, filtering through 70 μm Pre-Separation Filter (Miltenyi Biotec, 130-095-823) and centrifuging at 4°C (300 rcf, 5 minutes), cells were seeded on 24-well plate (0.5 x 10^5^ cell/well) in growth medium (DMEM medium; Thermo Fisher Scientific, 10569010, fetal bovine serum; VWR, 97068-085, 1% penicillin & streptomycin; Thermo Fisher Scientific, 15140122). The 24-well plates were collagen coated (Millipore Sigma, 125-50) before plating the cells. For siRNA treatment, *Tcf7l2* Silencer^®^ Select Pre-designed siRNA (Thermo Fisher Scientific, 4390771; s74838), *Ttll3* Silencer^®^ Select Pre-designed siRNA (Thermo Fisher Scientific, 4390771; s97693), and Silencer^®^ Select Negative Control #1 siRNA (Thermo Fisher Scientific, 4390843) were delivered by using Lipofectamine^®^ RNAiMAX Transfection Reagent (Thermo Fisher Scientific, 13778-075) according to the manufacturer’s instructions. The cells were treated with siRNA for 72 hours before further analysis.

### UCSC/JASPAR binding prediction and Cut and Run assay

UCSC/JASPAR bioinformatics database was used to search the promoter regions of individual genes in order to identify the binding profiles of transcription factors related to canonical Wnt signaling. Tcf7l2 predicted binding sites to the promoter region of *Ttll3* gene were identified at chr6:113388069-113388082. The following primers were used for the Cut and Run assay: forward TTGAACATCATGTGGCCCAGT and reverse CACTGTAAACCAGGGACTGCT. The soft palatal tissue from E13.5 embryos was collected and fixed according to the manufacturer’s instructions based on Cut and Run assay kit (Cell Signaling, 86652). Ten μl of IgG isotype control (Cell Signaling, 3900) or Tcf7l2 (Thermo Fisher Scientific, MA5-14935) were used for each reaction. Embryos from two litters were used for each replicate.

### CellChat and SCENIC analysis

Previously generated E13.5 soft palatal tissue single cell analysis data (Han et al. 2021) were processed using Seurat R package. RunPCA and RunUMAP visualization were used for cluster visualization. The data are available in GEO under accession number GSE155928. This dataset was processed with R package SCENIC to identify gene regulatory networks (Aibar et al. 2017). GENIE3 was used to identify the individual transcription factors that were later scored by AUCell. R package CellChat (Jin et al. 2021) was used to recognize the predicted ligand-receptor interactions between individual cell populations in the soft palate. The CellChat workflow was used with standard parameters (pre-processing functions: identifyOverExpressedGenes, identifyOverExpressedInteractions and projectData, core functions: computeCommunProb, computeCommunProbPathway and aggregateNet). netVisual_bubble and netAnalysis_signalingRole_heatmap were used to visualize the predicted results.

### Imaging

Immunofluorescence images were captured using a Leica DMI3000 B research microscope. Brightfield images were captured using a Keyence BZ-X710 system at the Center for Craniofacial Molecular Biology, University of Southern California.

### Statistical analysis

GraphPad Prism Software (Prism 9) was used for statistical analysis. We used N ≥ 3 for all experiments unless otherwise stated. For each qPCR experiment, at least three technical replicates were analyzed. For histological staining, immunofluorescence, and RNAScope, we used samples from at least three individual mice, or embryos, and representative images are shown. All bar graphs display mean ± s.e.m. Statistical comparisons were done using unpaired Student’s t-test with two-tailed calculations. One-way ANOVA was used for analyzing data for comparison of four independent groups. P < 0.05 was treated as statistically significant for all analyses.

## Data availability

The data for the RNA sequencing used in this study have been deposited in NCBI’s Gene Expression Omnibus (Edgar et al. 2002) and are accessible through GEO series accession number GSE208619. Previously generated E13.5 soft palatal tissue single cell analysis data (Han et al. 2021) and are available in GEO under accession number GSE155928.

## Acknowledgements

We are grateful to Bridget Samuels for critical reading of this manuscript and Giselle Mejia for the schematic illustrations. We acknowledge USC Libraries Bioinformatics Service for supporting with analysis of the sequencing data. The bioinformatics software and computing resources were funded by the USC Office of Research and the Norris Medical Library.

---

**Fig. S1.**
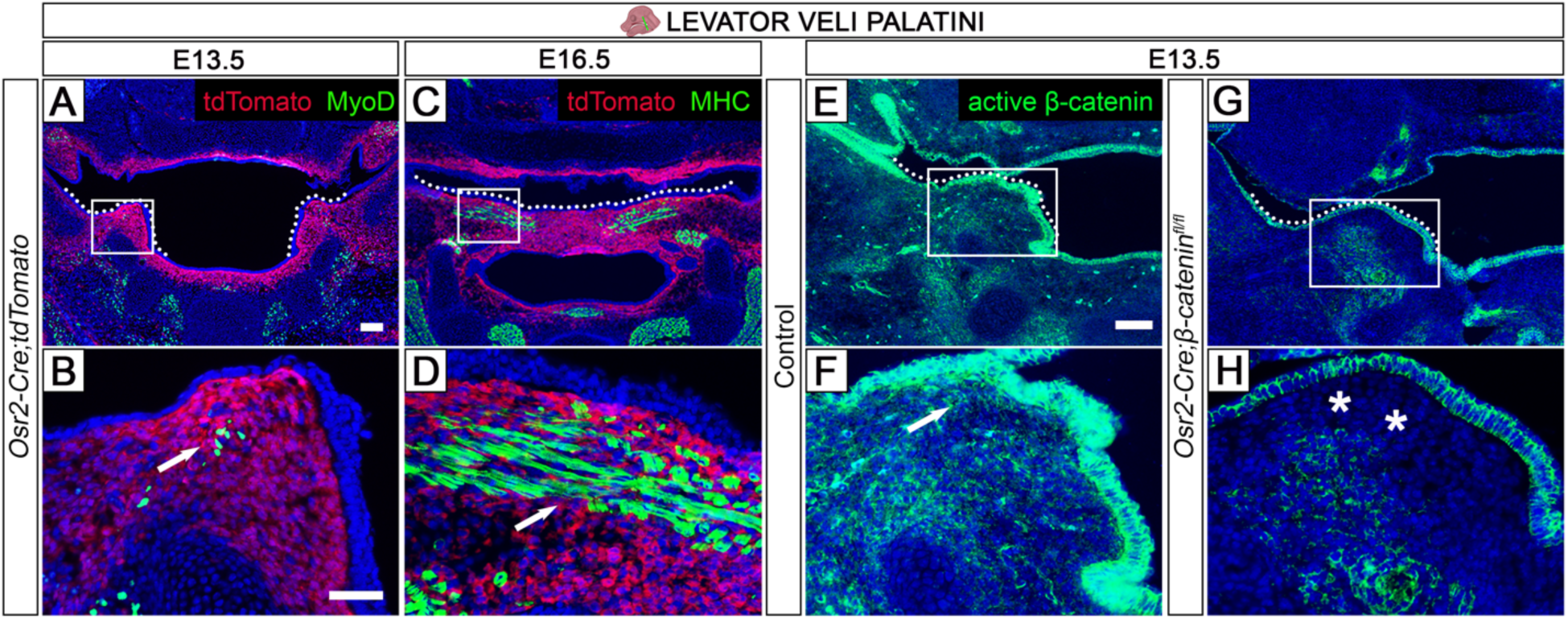
*Osr2-Cre* driver labels CNC-derived mesenchymal cells in the soft palate region. (A-D) *Osr2-Cre;tdTomato* mice at E13.5 (A-B) and E16.5 (C-D). *Osr2-Cre* driver targets only the palatal CNC-derived mesenchymal cells (red). The mesoderm-derived myogenic cells stained by MHC (green) are not targeted, which makes *Osr2-Cre* a highly useful tool for studying tissue-tissue interactions in the soft palate region. (E-H) Active β-catenin (green) immunofluorescence staining at E13.5 in control (E-F) and *Osr2-Cre;β-catenin^fl/fl^* mice (G-H). Boxes in A, C, E, and G show locations of higher magnification images in B, D, F, and H, respectively. Schematic drawing of the mouse head in the top panel depicts the position and angle of sectioning. White dotted lines indicate palatal shelves in A, C, E, and G. White arrows in B, D, and F point to positive areas. Asterisks in H show a lack of positive signal. Scale bar in A indicates 100 µm in A and C; scale bar in B indicates 50 µm in B, D, F, and H; scale bar in E indicates 100 µm in E and G.

**Fig. S2.**
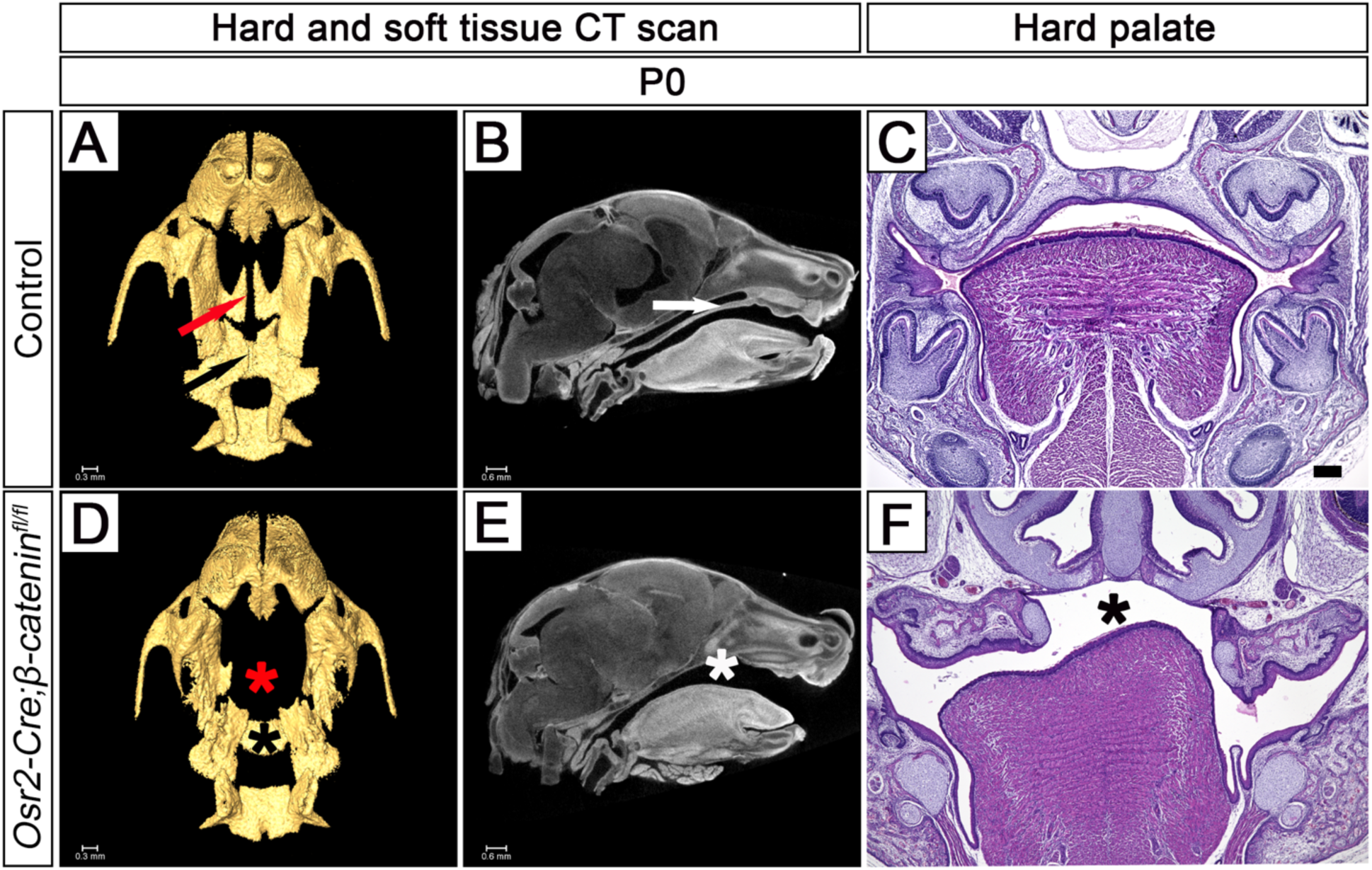
*Osr2-Cre;β-catenin^fl/fl^* mice display complete cleft palate. (A-B, D-E) Hard (A, D) and soft tissue (B, E) CT scans at P0 showing the defects of the palatine process of the maxilla (red asterisk in D), defects of the palatine bone (black asterisk in D), and absence of the palate in the *Osr2-Cre;β-catenin^fl/fl^*mice (white asterisk in E) in comparison to the normal palatine process of the maxilla (red arrow in A), palatine bone (black arrow in A), and hard palate development (white arrow in B) in controls. (C, F) Hematoxylin and eosin staining at the level of the hard palate in control (C) and *Osr2-Cre;β-catenin^fl/fl^* mice (F) at P0. Black asterisk in F identifies the cleft palate in *Osr2-Cre;β-catenin^fl/fl^*. Scale bar in A indicates 0.3 mm in A, and D; scale bar in B indicates 0.6 mm in B, and E; scale bar in C indicates 200 µm in C and F, respectively.

**Fig. S3.**
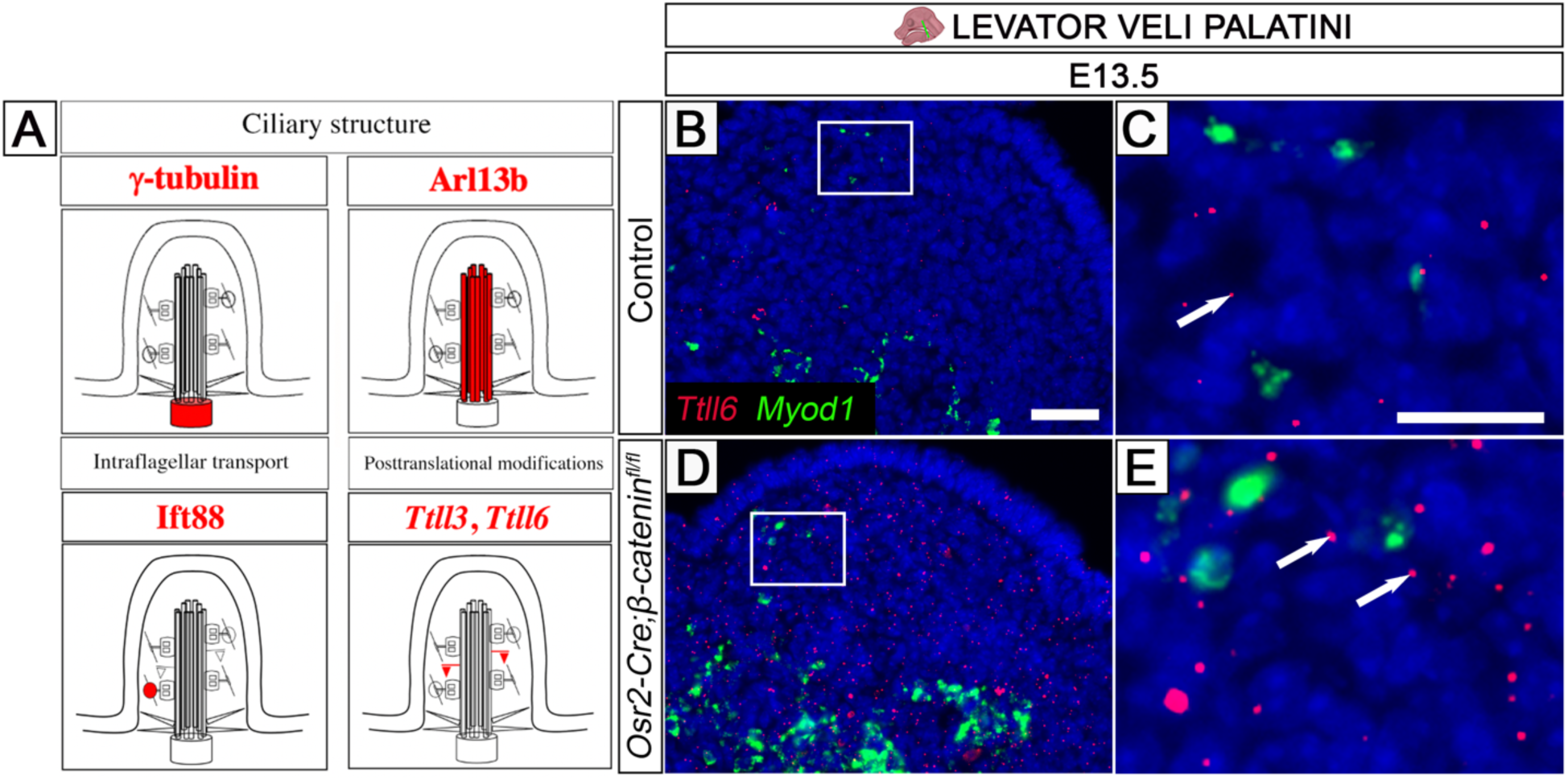
Description of the ciliary phenotype. (A) Schematic drawings of primary cilia focusing on ciliary structure (basal body and ciliary axoneme), intraflagellar transport and posttranslational modifications. (B-E) *Ttll6* (red) and *Myod1* (green) *in situ* RNAScope hybridization in control (B-C) and *Osr2-Cre;β-catenin^fl/fl^*mice (D-E). Schematic drawing of the mouse head in the top panel depicts the position and angle of sectioning. Boxes in B and D show approximate location of images in C and E. White arrows in C and E point to positive signals. Scale bar in B indicates 50 µm in B, and D; scale bar in C indicates 20 µm in C and E.

